# Distributed theta networks support the control of working memory: Evidence from scalp and intracranial EEG

**DOI:** 10.1101/2025.08.14.670214

**Authors:** Lingxiao Shi, Kaustav Chattopadhyay, Samantha M. Gray, Josset B. Yarbrough, David King-Stephens, Ignacio Saez, Fady Girgis, Ammar Shaikhouni, Stephan U. Schuele, Joshua M. Rosenow, Eishi Asano, Olivia Kim McManus, Shifteh Sattar, Robert T. Knight, Elizabeth L. Johnson

**Author notes:** **Conflicts of interest:** None.

## Abstract

We combined scalp EEG and intracranial EEG (iEEG) to identify spectral and network-level signatures of executive control during a delayed match-to-sample task working memory task. To isolate executive processes, we contrasted test and sample phases, matched in perceptual input but differing in cognitive demand. Scalp EEG revealed increased frontal midline theta event-related spectral perturbations (ERSPs), dynamic increases and decreases in posterior theta-alpha ERSPs, and decreased central alpha-beta ERSPs during the test phase. These local spectral changes were accompanied by enhanced frontoposterior theta phase synchrony and network hub strength, predicting higher behavioral accuracy. Using a novel cross-modal scalp EEG-iEEG ERSP similarity approach, we localized the sources of scalp-derived frontal midline, posterior, and central control effects to medial frontal, parietal, temporal, and occipital regions. Our results integrate power and connectivity measures across scalp and iEEG, linking local spectral fluctuations to broader network organization. Together, they support a model in which executive control emerges from flexible, temporally precise coordination between medial frontal control hubs and posterior representational systems.

## Introduction

Working memory (WM) is a limited-capacity system supporting the temporary storage and manipulation of information essential for goal-directed behavior. A core function of WM is coordinating incoming sensory input with retrieved long-term memories, a process dependent on executive control (Baddeley, 2000).

According to the multicomponent model of WM, sensory modality-specific storage subsystems such as the phonological loop, visuospatial sketchpad, and episodic buffer are regulated by a modality-independent central executive that governs attention, resolves interference, and updates representations based on task demands (Baddeley & Hitch, 1974; Baddeley, 1996, 2000). These theoretical accounts have sparked a growing interest in the brain mechanisms that support executive control in WM, especially those that coordinate and manipulate information over time, a role often linked to the frontal lobe (Collette & Van der Linden, 2002; D’Esposito et al., 1995; Smith & Jonides, 1999; Johnson et al., 2017, 2022).

Frontal midline theta (FMT; 4-8 Hz) activity is linked to executive control during maintenance and manipulation of information over short time scales (Cavanagh & Frank, 2014; Cavanagh & Shackman, 2015; Hsieh & Ranganath, 2014; Itthipuripat et al., 2013; Sauseng et al., 2010; Maurer et al., 2015; Mitchell et al., 2008). Electrophysiological studies across EEG, MEG, and intracranial recordings demonstrate that FMT power increases with WM load (Gevins et al., 1997; Jensen & Tesche, 2002; Meltzer et al., 2008; Muthukrishnan et al., 2020). FMT is also implicated in the maintenance of temporal order: greater FMT power emerges when participants must preserve the sequence of items rather than their identity alone (Hsieh et al., 2011; Roberts et al., 2013). Together, these findings suggest that FMT reflects a domain-general mechanism for sustaining goal-relevant representations and for temporally structured information.

Posterior cortical regions, particularly sensory and parietal areas (Collette et al., 1999), generate alpha (8-14 Hz) and beta (13-30 Hz) rhythms during WM maintenance (Erickson et al., 2017). Alpha power varies across WM phases: during encoding, occipitotemporal alpha suppression reflects enhanced visual processing and improved performance under high memory load (Bonnefond & Jensen, 2012; Wianda & Ross, 2019).

During retention, alpha power rebounds, inhibiting irrelevant sensory inputs and protecting WM representations from interference (Bonnefond & Jensen, 2012; Wianda & Ross, 2019). This dual role is supported by cross-frequency coupling (CFC), where alpha phase modulates gamma (30-50 Hz) power to organize local neural activity and maintain WM content (Axmacher et al., 2010; Wianda & Ross, 2019; Davoudi et al., 2021).

Interregional alpha-band synchrony, measured by phase-locking value (PLV), further predicts spatial coding and performance in WM (Norouzi & Daliri, 2024). Beta rhythms are proposed to stabilize WM representations and clear information during updating (Schmidt et al., 2019). Beta bursts have also been linked to state transitions that interrupt default processing (Lundqvist et al., 2016), and impaired alpha-beta modulation during WM tasks has been observed in schizophrenia (Adams et al., 2020; Moran et al., 2018). Together, these findings underscore alpha and beta rhythms as key mechanisms for balancing sensory gating, interference suppression, and the dynamic updating of WM.

WM function depends on local neural activity, interregional phase synchrony, and cross-frequency interactions. Theta rhythms are critical for distributed network integration, while high-frequency broadband activity facilitates local computations (Axmacher et al., 2010; Canolty et al., 2006; Jensen & Lisman, 2005). FMT coordinates interactions between frontal and posterior cortices via long-range theta and theta-alpha phase synchrony (Fell & Axmacher, 2011; Liebe et al., 2012; von Stein & Sarnthein, 2000). This frontoparietal synchrony, often measured by PLV and graph-theoretic metrics, supports the integration of executive and sensory processes and scales with WM performance (Albouy et al., 2017; Palva et al., 2010; Sauseng et al., 2005; Shin et al., 2022). Conversely, disruptions in this network have been linked to cognitive deficits in schizophrenia and age-related decline (Griesmayr et al., 2014; Reinhardt & Nguyen, 2019). Computational and modeling studies further suggest that theta–gamma CFC provides a temporal scaffold for sequencing WM content (Axmacher et al., 2010; Jensen & Lisman, 2005; Lisman & Jensen, 2013; Siegel et al., 2009). Together, these findings underscore the role of theta and alpha rhythms in organizing and synchronizing distributed brain activity to support the temporal precision and flexibility of WM.

Non-invasive brain stimulation techniques underscore the causal role of theta rhythms in WM. In-phase theta transcranial alternating current stimulation (tACS) enhances WM retrieval performance when applied to bilateral prefrontal (PFC) or parietal regions during WM encoding (Jaušovec & Jaušovec, 2014; Hosseinian et al., 2021; Tseng et al., 2018; Debnath et al., 2025), and anti-phase theta tACS interferes with endogenous rhythms and impairs WM (Chander et al., 2016; Tseng et al., 2018; Alekseichuk et al., 2017). Transcranial direct current stimulation (tDCS) also enhances frontoparietal theta synchrony and theta-gamma CFC, both of which are associated with improved WM performance (Hu et al., 2022; Johnson et al., 2022; Jones et al., 2017, 2020). Frequency band-limited transcranial magnetic stimulation (TMS) provides additional evidence for frequency- and region-specific roles in WM: frontal theta TMS facilitates prioritization of information, while parietal alpha TMS supports suppression of irrelevant inputs (Li et al., 2017; Riddle et al., 2020). TMS-EEG studies further demonstrate that rhythmic stimulation can entrain long-range synchrony, including evidence for bottom-up theta network dynamics from posterior sensory to frontal areas (Chung et al., 2018; Li et al., 2017; Miyauchi et al., 2016).

Lesion studies provide converging evidence for the roles of the PFC and parietal cortex in theta-band coordination and WM functions. Tasks requiring spatiotemporal integration and executive control rely on intact lateral PFC function (Johnson et al., 2017; Parto Dezfouli et al., 2021; Davoudi et al., 2021; Postle et al., 1999), and attentional maintenance of mnemonic representations relies on intact parietal function (Berryhill et al., 2011). Healthy individuals exhibit bidirectional fronto-posterior interactions during WM, with frontal low-theta supporting demand-sensitive executive control and posterior alpha-beta providing a PFC-independent pathway that can sustain performance when frontal contributions are reduced (Johnson et al., 2017). Patients with lateral PFC damage exhibit attenuated slow-theta activity, reduced frontoposterior connectivity, and disruptions in frontal and mesial posterior signatures of spatiotemporal integration (Parto Dezfouli et al., 2021; Davoudi et al., 2021). Together, these findings highlight the PFC’s essential role in orchestrating large-scale oscillatory dynamics that bind spatial and temporal information into coherent, goal-relevant representations.

We investigated the spectral patterns underlying sequence maintenance and comparison using scalp EEG and intracranial EEG (iEEG) using a delayed match-to-sample (DMS) task (Cross et al., 2025; Yarbrough et al., 2025; **Fig. 1A**). On each trial, participants encode a sample sequence of three geometric shapes (varying in color, shape, and location) during a study phase, maintain the sequence across a delay, and compare the sample to a test sequence where one dimension may differ. By isolating the comparison process, we test three hypotheses: (1) FMT power increases while posterior alpha power decreases during the test relative to sample sequence indexing executive demand and distractor inhibition. (2) Interregional FM-posterior theta PLV increases during the test relative to the sample sequence. (3) FMT and posterior theta network connectivity, quantified by node degree based on PLV, increase, measuring enhanced network connectivity. To test these hypotheses, we examined changes in event-related spectral perturbations (ERSPs), phase synchrony between brain regions (PLV), and overall network connectivity using graph theory. We further incorporate iEEG data to provide putative information on the spatial sources of scalp EEG-derived ERSPs.

**Figure 1.**
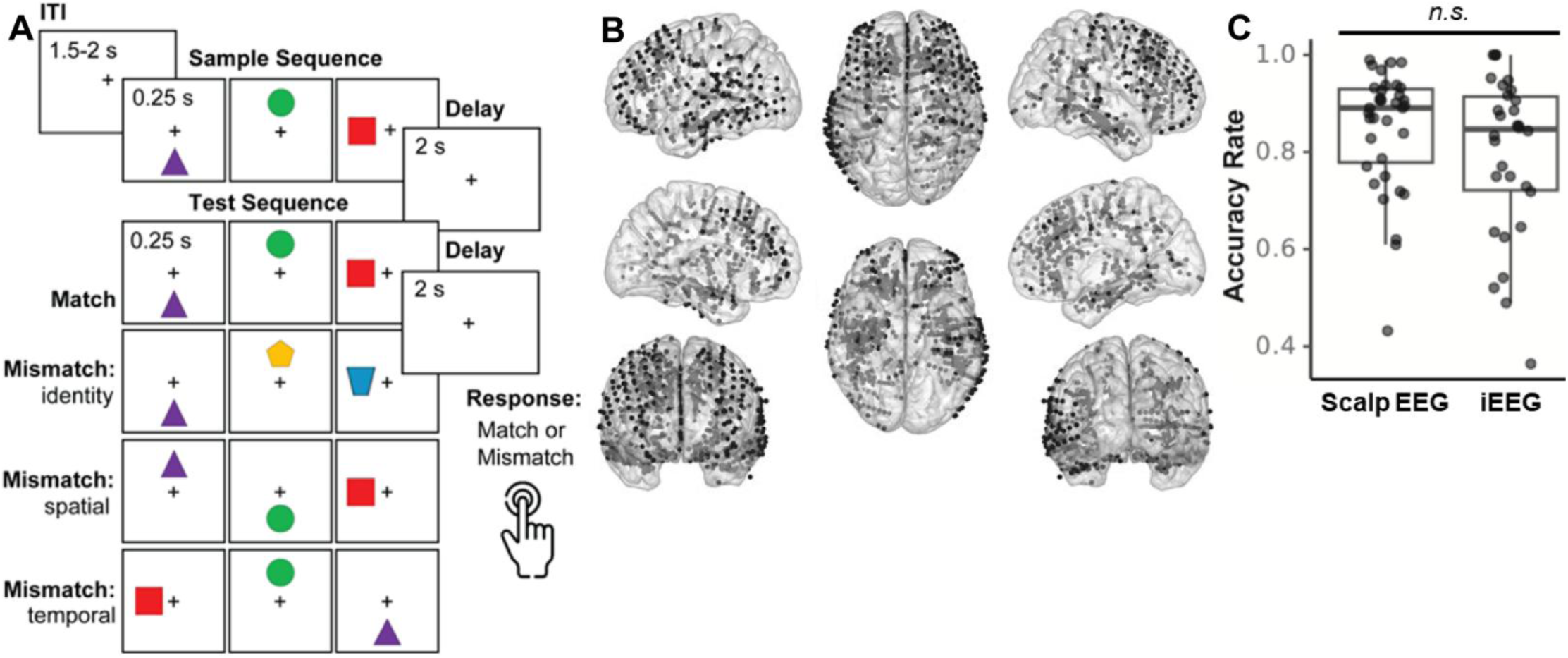
WM-DMS task schematic, iEEG electrode coverage, and task accuracy across modalities. **A)** In each trial, participants viewed a sample sequence of three geometric shapes (0.25 s stimulus, 0.25 s interstimulus interval/ISI [not shown]), followed by a 2-s delay and a test sequence with identical timing. The test either matched the sample or differed in one dimension. Shown are four trial types: exact match, identity mismatch, spatial mismatch, and temporal order mismatch. ITI = intertrial interval. **B)** Spatial distribution of seizure- and artifact-free bipolar iEEG channels (n = 1,315) from all patients (n = 29) shown in MNI space. Each black dot marks the midpoint between a bipolar electrode pair for visualization purposes. **C)** Boxplots of behavioral accuracy for scalp EEG (n = 35) and iEEG (n = 29) participants show comparable performance across modalities. Black dots represent individual data points.

## Methods

### Working memory delayed match-to-sample task

Participants performed our WM-DMS task, in which they detected mismatches in sequences of three colored shapes (see **Fig. 1A**). On each trial, they maintained central fixation for 1.5-2.0 s (i.e., jittered intertrial interval). Then, they viewed a sample sequence of three different shapes in specific spatiotemporal positions (0.25 s stimulus, 0.25 s ISI). Each shape appeared in one of four different spatial positions (left, right, top, bottom), followed by a 2-s delay and one of four conditions of test sequence with identical timing to the sample sequence. The test sequence matched the sample sequence exactly (i.e., match) or was changed on one dimension: two out of three stimuli were either new (mismatch: identity), swapped across spatial positions (mismatch: spatial), or swapped across temporal positions (mismatch: temporal). On mismatch trials, one stimulus always matched on all three dimensions, and the temporal positions of the two mismatched stimuli were evenly counterbalanced (i.e., 1 and 2, 2 and 3, 1 and 3). Following a second 2-s delay, they responded match/mismatch/unsure by self-paced mouse click. The central fixation crosshair remained on screen for the duration of the task. Following eight practice trials, participants completed 256 (scalp EEG) or 128 (iEEG) trials with a break every 16 trials. The four test conditions were evenly counterbalanced and presented in pseudo-randomized order with four trials per condition every 16 trials. For half of all trials, a star flashed for 0.1 s midway through both delays at randomly jittered times ranging from 1.0-1.15 s from the onset of the delay. The star trials were evenly counterbalanced across conditions and appeared on eight of every 16 trials. The other half of trials were considered control trials and included in EEG analysis in this study. Behavioral data were collected using custom-built MATLAB (MathWorks Inc., Natick, MA) scripts with the Psychtoolbox-3 software extension.

Behavioral accuracy rate was calculated per participant as the hit rate (i.e., proportion of match sequences that were correctly identified as match) minus false alarm rate (proportion of mismatched sequences that were incorrectly identified as match) (Snodgrass & Corwin, 1988). This procedure defines chance accuracy as zero and corrects for differences in trial counts between conditions as well as an individual’s tendency to respond match/mismatch. Response time (RT) was calculated per participant across all trials.

### Scalp EEG

#### Participants

Participants were 35 cognitively healthy adults (20 females; M ± SD, age: 25.26 ± 7.02 years; education: 15.15 ± 2.28 years) with normal/corrected-to-normal vision and hearing, and no neurological or psychiatric diagnoses. Sensitivity analysis indicated that this sample size provides 80% power to detect small-medium effects at two-tailed α < 0.05 (dependent-samples t-test Cohen’s d = 0.49 (Faul et al., 2007); correlation ρ = 0.44 (Erdfelder et al., 2009)). Participants were included based on above-chance task performance (inclusion criterion: <0.35 error in each condition, chance 0.5 (Johnson et al., 2023)). All participants provided written informed consent in accordance with the Declaration of Helsinki as part of the research protocol approved by the University of California, Berkeley Institutional Review Board.

#### Data acquisition and preprocessing

EEG data were recorded at a sampling rate of 1,024 Hz using a 64 + 8 channel BioSemi ActiveTwo amplifier with Ag-AgCl pin-type active electrode channels mounted on a cap based on the extended 10-20 system (BioSemi, Amsterdam, NL). The horizontal electrooculogram (EOG) was recorded at both external canthi, and the vertical EOG was monitored with a right inferior eye channel. Two channels were placed on the earlobes for offline referencing. Impedances were kept below 20 kΩ. Raw data were bandpass-filtered with a 0.1-100-Hz finite impulse response filter, and 60-Hz line noise was removed using the discrete Fourier transform. Filtered data were down-sampled to 512 Hz, demeaned, and epoched into 8.5-s trials (−1 s from the onset of the sample sequence to +1 s from the offset of the post-test delay). Data were inspected blind to channel locations and experimental task parameters, and any channels displaying artifactual signal (from poor contact, machine noise, etc.) were excluded. Independent component analysis was performed on the remaining channels to remove EOG, electromyography, and other artifacts (Hipp & Siegel, 2013). Any channels that had been removed were replaced with the interpolated mean values from neighboring channels (M = 7.6 channels), and data were manually re-inspected to remove trials containing residual noise. The surface Laplacian spatial filter was then applied to minimize volume conduction and enhance the source signal (Cohen, 2015; Lai et al., 2018). Trials with a flashing star were excluded from EEG analysis. The event-related potential (ERP) was computed across all remaining control trials and subtracted from each trial to isolate induced, non-phase-locked activity related to the task (Johnson et al., 2017, 2018, 2019, 2022; Jones, Johnson, & Berryhill, 2020; Jones, Johnson, Tauxe, et al., 2020). Finally, error trials were excluded (Davoudi et al., 2021; Johnson et al., 2017, 2018, 2019, 2023; Parto Dezfouli et al., 2021), resulting in 101.57 ± 14.53 (M ± SD) artifact-free, correct trials analyzed per participant. Preprocessing and analysis routines utilized functions from the open-source FieldTrip toolbox for MATLAB (Oostenveld et al., 2011).

#### Spectral decomposition

The preprocessed data segments were zero-padded to the next power of 2 to minimize filtering-induced edge artifacts and the Hanning taper time-frequency spectrum was calculated by sliding a frequency-dependent window of three cycles (Δt = 3/f) in 50-ms steps at each frequency from 2.5-49.5 Hz (1-Hz resolution, 2-Hz bandwidth). Power outputs were extracted for the pre-stimulus baseline (−0.75 to −0.25 s from sample onset) and post-stimulus time series (−0.25-6.5 s from sample onset), and relative-change baseline-corrected (i.e., (post-stimulus time series – baseline mean)/baseline mean). The resulting ERSPs were averaged across trials per participant for group-level statistical analysis.

#### Functional connectivity

Spectral decomposition was performed on a subset of 2.5-20.5 Hz frequencies using the same procedures, and phase outputs were extracted for the post-stimulus time series (−0.25-6.5 s from sample onset). Functional connectivity was quantified per timepoint as the PLV independent of amplitude (Lachaux et al., 1999). This method calculates the consistency in channel-pair phase differences across a series of data points, here, trials per timepoint. PLV was computed across all channel pairs. In addition to analyzing inter-channel PLV, we applied graph theory to map the topographical distributions of PLV networks over time. According to graph theory, brain networks are collections of nodes (here, Laplacian-transformed channels) and inter-node edges (PLV), summarized as adjacency matrices (Sporns, 2018). To define PLV adjacency matrices, PLV outputs were assessed for network degrees (i.e., the weight of connections between each channel and all other channels) using an individually defined threshold of 0.65 relative to each participant’s maximum PLV (Johnson et al., 2022; Jones, Johnson, Tauxe, et al., 2020).

#### Statistics

Spectral WM effects were identified using dependent-samples t-tests with cluster-based correction for multiple comparisons (Maris & Oostenveld, 2007). Post-stimulus data were separated into sample (0-3.25 s from sample onset) and ‘test’ time series (3.25-6.5 s from sample onset, i.e., 0-3.25 s from delayed match onset; see **Fig. 1A**), and the test time axis was re-coded as 0-3.25 s to match the sample time axis. Clusters were formed in space, time, and frequency by thresholding sample vs. test t-statistics at p < 0.05 using the maximum sum criterion. Spatial adjacency was defined by the BioSemi-64 template and clustering by a typical minimum of two channels. Permutation distributions were generated by randomly shuffling sample and test data labels (1,000 iterations), and corrected p-values were obtained by comparing the observed data to the random permutation distributions. This approach recreates any biases in the data with each randomization and tests for effects against the global null hypothesis without any assumptions about their spatial, temporal, or spectral distribution. Cluster-based statistics were implemented using the FieldTrip toolbox for MATLAB (Oostenveld et al., 2011).

Post-hoc correlations examined relationships between spectral effects (e.g., ERSP and PLV effects) to quantify associations between regional activity and interregional network effects, and between spectral effects and individual behavioral outcomes (i.e., accuracy rate, RT). Spearman’s rank correlation coefficients were calculated from spectral sample versus test differences averaged across cluster-corrected data points (Johnson et al., 2022).

### Intracranial EEG

#### Participants

Participants were 29 cognitively healthy adults (8 females; M ± SD, age: 29.20 ± 10.75 years) with normal/corrected-to-normal vision and hearing, who were undergoing intracranial monitoring as part of clinical management of seizures. Participant ages did not differ from the scalp EEG dataset (BF_10_ < 1). The initial sample included 30 patients selected based on above-chance task performance (inclusion criterion: <0.35 error in each condition, chance 0.5 (Johnson et al., 2023)). One patient was excluded due to cortical atrophy across the implanted hemisphere. Neurosurgical patients were recruited from the University of California, Irvine, the University of California, Davis, the University of California, San Francisco, Mt. Sinai hospitals, California Pacific Medical Center, Nationwide Children’s Hospital, Northwestern Memorial Hospital, Children’s Hospital of Michigan, and the University of California, San Diego Rady Children’s Hospital. All participants provided written informed consent in accordance with the Declaration of Helsinki as part of the research protocol approved by the Institutional Review Board at each hospital.

#### Channel placement and localization

Macro-electrodes were surgically implanted for extra-operative recording based solely on the clinical needs of each patient. These electrode channels were placed subdurally in grids and strips with 10-mm spacing (i.e., electrocorticography/ECoG) and/or stereotactically in tracks with 3-5mm spacing (stereotactic EEG/sEEG). Anatomical locations were determined by co-registering post-implantation computed tomography (CT) coordinates to pre-operative magnetic resonance (MR) images, as implemented in FieldTrip (Stolk et al., 2018). Channels were localized in native space based on visual inspection of individual anatomy and transformed into standard MNI space for group-level analysis. BrainNet Viewer was used to visualize channels on the MNI-152 template brain (Xia et al., 2013) (see **Fig. 1B**).

#### Data acquisition and preprocessing

iEEG data were recorded at a sampling rate of 200-5,000 Hz using Nihon Kohden and Natus recording systems, and data sampled >1,000 Hz were resampled to 1,000 Hz offline. As in the analysis of scalp EEG data, spectral decomposition was performed up to 50 Hz, and so the lowest sampling rate of 200 Hz is double the minimum Nyquist frequency required for analysis (i.e., 2 cycles/frequency = 100 Hz). Raw data were filtered with 0.1-Hz high-pass and, if sampled at 1,000 Hz, 300-Hz low-pass finite impulse response filters, and 60-Hz line noise harmonics were removed using discrete Fourier transform. Filtered data were demeaned and epoched into 8.5-s trials (−1 s from the onset of the sample sequence to +1 s from the offset of the post-test delay). Data were inspected blind to channel locations and experimental task parameters, and channels overlying seizure onset zones and any channels and epochs displaying epileptiform activity or artifactual signal (from poor contact, machine noise, etc.) were excluded, ensuring that data considered for analysis represent healthy tissue (Rossini et al., 2017). Neighboring channels within the same anatomical structure were then bipolar-referenced using consistent conventions: for ECoG, the subtraction was performed in the anterior-to-posterior direction; for sEEG, the subtraction was performed from deep to surface contacts (Johnson et al., 2018, 2019, 2023). ECoG grid channels were referenced on a row-by-row basis. A channel was discarded if it did not have an adjacent neighbor, if its neighbor was in a different anatomical structure, or, in the case of sEEG, if both the channel and its neighbor were in white matter. Bipolar montages were used to minimize contamination from volume conduction (Shirhatti et al., 2016), comparable to the Laplacian spatial filter used on the scalp EEG data (Mercier et al., 2022). Referenced data were manually re-inspected to remove trials containing residual noise. As described above for scalp EEG data, trials with a flashing star were excluded, the ERP was computed across all remaining control trials and subtracted from each trial, and then error trials were excluded, resulting in 51.66 ± 9.79 (M ± SD) seizure- and artifact-free, correct trials analyzed per participant.

#### Spectral decomposition and statistics

The Hanning taper time-frequency spectrum was calculated from the preprocessed data, as described above for scalp EEG data. The output resolution of 50 ms equated the ERSP time axis across iEEG datasets with different sampling rates and matched that of the scalp EEG ERSPs. Spectral decomposition procedures, time-frequency axes, and sample sizes were equated across iEEG and scalp EEG ERSPs because these input data characteristics affect cluster-based permutation test outputs (Sassenhagen & Draschkow, 2019). To achieve stable, power-controlled ERSPs that were standardized across participants, we applied bootstrapping to transform individual iEEG ERSPs for group-level analysis of trial-averaged data with the same statistical power as the scalp EEG data. Specifically, we randomly selected (without replacement) and averaged five trials per participant to create a new ‘trial’. We repeated this step 35 times per participant and then concatenated channels from all participants onto a single MNI brain for group-level statistical analysis (Oehrn, 2023). ERSP WM effects were analyzed using a dependent-samples t-test with cluster-based correction for multiple comparisons, as described above for scalp EEG data. Spatial adjacency was defined by a 15 mm distance and clustering without a minimum number of channels to permit both spatial clustering and identification of localized effects.

#### Similarity analysis

Similarity analysis identified iEEG channels exhibiting significant ERSP WM effects with time-frequency representations that were most like significant scalp EEG ERSP WM effects. Similarity analysis directly compares spatial patterns of neural activity across modalities to identify shared functional organization (Kriegeskorte et al., 2008; Cichy et al., 2014). Unlike forward modeling, which predicts scalp potentials from iEEG sources using head conductivity models (He et al., 2018), or inverse modeling, which reconstructs sources from scalp EEG via ill-posed estimation (e.g., sLORETA, beamforming), similarity analysis is entirely empirical as it requires no assumptions about tissue conductivity, source extent, or orientation. By using measured iEEG spectral signatures as a reference, this approach enables direct EEG-iEEG structural mapping, offering complementary anatomical specificity to model-based methods when sufficient intracranial coverage is available.

First, three channel-averaged references were defined by spatially clustered scalp EEG effects, yielding FM (channels AF3, AFz, AF4, Fp1, Fpz, Fp2, Fz), central (FC3, FC2, C1, C3, CP3, CP1, CPz, FCz, Cz), and bilateral posterior (P4, P6, P8, P10, PO4, PO8, P3, P5, P7, P9, PO3, PO7) references of statistically significant effects (i.e., masked time-frequency representation of t-statistics). Then, the time-frequency representation of statistically significant effects in each iEEG channel across all frequencies from 2.5-49.5 Hz, identical to scalp ERSP analysis (i.e., masked time-frequency representation of t-statistics), was analyzed point-by-point for similarity to each of the three scalp EEG references. The significant effects in all scalp EEG reference- and iEEG channel-masked time-frequency representations were binarized as 1 for positive effects and −1 for negative effects and analyzed by means of a similarity index as described above. For each time-frequency point with a significant effect in both representations, if the effect matched (i.e., both positive or negative), the index was given a similarity of 1, and if the effect did not match (i.e., one positive and one negative), the index was given a similarity of −1. The time-frequency representation of similarity for each scalp EEG-iEEG comparison was summed and normalized by dividing the sum by the number of datapoints compared (i.e., number of time-frequency points with a significant effect in both the scalp EEG and iEEG representations). The resulting similarity indices are bounded from −1 to 1, with −1 indicating perfect dissimilarity, 0 indicating no similarity, and 1 indicating perfect similarity.

Statistical analysis of similarity was performed using statistical bootstrapping. For each of the three reference images, we randomly selected (without replacement) and averaged the similarity indices of five iEEG channels. We repeated this step 1,000 times to create bootstrapped distributions of ERSP similarity data.

Similarity indices were z-scored on the bootstrapped distributions, and iEEG channels with similarity above threshold (z = 1.96, equivalent to two-tailed α = 0.05) were considered probable sources of scalp EEG effects. iEEG channels with no significant ERSP effects were excluded from analysis.

### Data and code availability

De-identified data and custom-built MATLAB codes will be available on OSF upon acceptance.

## Results

### Similar task performance in scalp EEG and iEEG samples

Behavioral accuracy rates did not differ between the scalp EEG (M ± SD: 0.848 ± 0.124; one-sample t(34) = 40.353, p < 0.001, Cohen’s d = 6.821) and iEEG (0.792 ± 0.167; one-sample t(28) = 25.531, p < 0.001, Cohen’s d = 4.741) samples (BF_10_ < 1; **Fig. 1C**). Significant negative correlations between accuracy and RT were observed in both the scalp EEG (ρ = −0.340, p = 0.046) and iEEG (ρ = −0.425, p = 0.022) samples, and did not differ between samples (z = 0.380, p = 0.704), with individuals with superior performance responding faster. These results provide converging evidence that WM-DMS performance was similar across samples— i.e., that neurosurgical patient performance was consistent with that of our non-clinical sample.

### Theta signature of test sequence encoding and beta signature of sample sequence encoding

To identify spectral signatures of WM, we compared the sample and test sequence scalp EEG ERSPs from the onset of the first stimulus to the offset of the delay. A significant cluster-corrected t-test revealed test > sample and sample > test ERSP effects that varied in time and frequency across the scalp topography (**Fig. 2A-C**). Sample > test effects were characterized by stimulus onset-locked power in the theta-beta range (5-35 Hz) in bilateral posterior channels and stimulus offset-locked power focused in the beta range (15-35 Hz) in frontal and central channels (p = 0.0001, Cohen’s d = 2.539). By contrast, test > sample effects were characterized by stimulus offset-locked power focused in the theta range (3-10 Hz) in all channels (p = 0.009, Cohen’s d = 1.687) as well as beta range (15-30 Hz) in bilateral posterior channels (p = 0.045, Cohen’s d = 1.275). Additional sample > test effects were characterized by sustained power in the alpha-beta range (10-30 Hz) during the delay in central channels (p = 0.025, Cohen’s d = 0.556). These results demonstrate a FM beta ERSP signature of sample sequence encoding and theta ERSP signature of test sequence encoding (see **Fig. 2A**), a bilateral posterior theta-beta ERSP signature showing temporally alternating engagement during sample and test sequence encoding (see **Fig. 2B**), and a central alpha-beta ERSP signature of sample sequence maintenance (see **Fig. 2C**). Correlations between ERSP effects and behavioral outcomes did not reach significance (p > 0.12).

**Figure 2.**
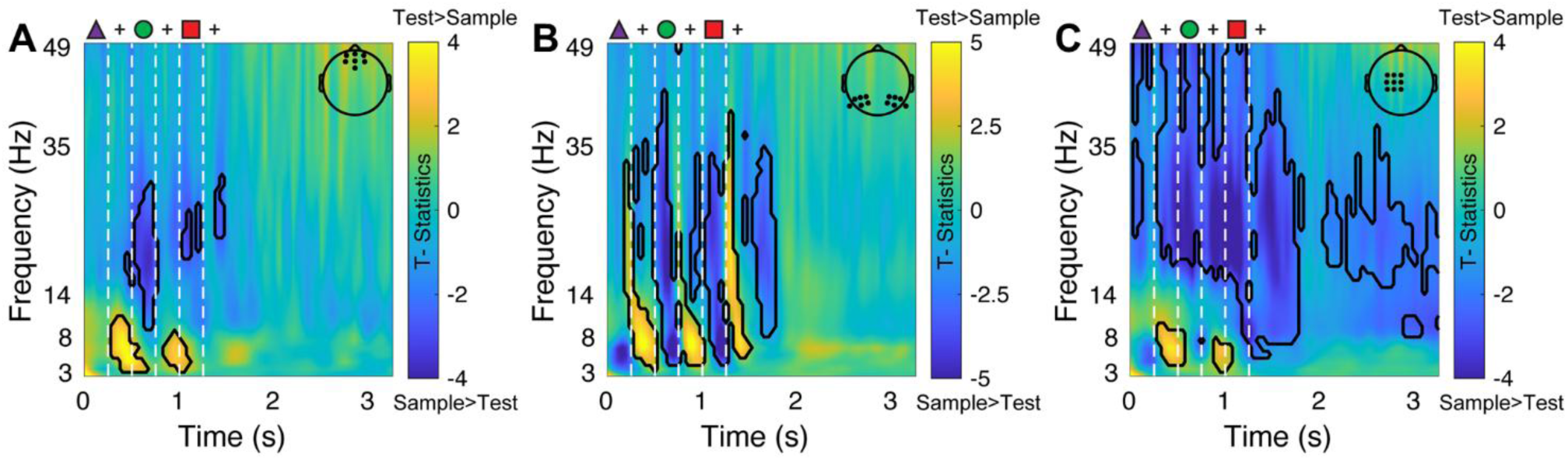
Task-related scalp EEG ERSP effects. Time-frequency representations of t-statistics on power differences (test vs. sample) across FM, bilateral posterior, and central scalp EEG channels (insets). Time zero marks the onset of the first stimulus in each sequence. Geometric symbols indicate the three stimulus onsets (triangle, circle, square), and the plus sign denotes ISIs and delays. **A)** FM channels show increased theta ERSPs and decreased beta ERSPs during test sequence encoding (yellow = test > sample; blue = sample > test). **B)** Bilateral posterior channels show increased theta-beta ERSPs during ISIs and reduced theta-beta ERSPs during stimulus presentation during test sequence encoding. **C)** Central channels show increased theta ERSP during test sequence encoding and sustained alpha-beta suppression from test sequence onset through the delay. Black contours outline significant clusters.

### Spectral signatures of working memory across broadly distributed neuroanatomy

To infer the likely neuroanatomical sources of scalp EEG effects, we computed stable, power-controlled iEEG ERSPs that were standardized across participants, concatenated the iEEG ERSPs from all channels onto a single MNI brain (Oehrn, 2023), and performed group-level statistical analysis of iEEG spectral signatures of WM. This analysis yielded cluster-corrected ERSP effects in 1,305/1,315 iEEG channels (p < 0.05; see **Fig. 1B**). One-sample t-tests revealed that, across all channels, iEEG effects were similar to FM (M ± SD = 0.082 ± 0.364; p = 8×10^-16^), bilateral posterior (0.057 ± 0.269; p = 4×10^-14^), and central (0.047 ± 0.333; p = 4×10^-7^) scalp EEG effects. Additionally, paired-samples t-tests revealed that iEEG effects were more similar to FM than bilateral posterior (p = 4×10^-7^) and central (p = 3×10^-15^) scalp EEG effects. The difference in the similarity of iEEG effects to bilateral posterior versus central scalp EEG effects did not reach significance (p = 0.054). Statistical bootstrapping identified the iEEG channels exhibiting significant ERSP WM effects that were either similar (z > 1.96, p < 0.05) or dissimilar (z < −1.96, p < 0.05) to significant FM, bilateral posterior, and central scalp EEG ERSP WM effects, respectively.

Analysis of similarity to FM scalp EEG effects (see **Fig. 2A**) revealed 284 (107 unique) iEEG channels with similar spectral signatures and 236 (96 unique) iEEG channels with dissimilar spectral signatures (all p < 0.05). Similar spectral signatures were distributed across rostral-superior and medial frontal regions, as well as temporal and occipital regions (**Fig. 3A-B**). Dissimilar spectral signatures were distributed across lateral PFC regions (**Fig. 3C**). Analysis of similarity to bilateral posterior scalp EEG effects (see **Fig. 2B**) revealed 265 (100 unique) iEEG channels with similar spectral signatures and 239 (96 unique) iEEG channels with dissimilar spectral signatures (all p < 0.05). Similar spectral signatures were distributed across inferior parietal, temporal, and occipital regions, as well as medial frontal regions (**Fig. 4A-B**). Dissimilar spectral signatures were distributed across lateral PFC and medial sensorimotor regions (**Fig. 4C**). Analysis of similarity to central scalp EEG effects (see **Fig. 2C**) revealed 277 (88 unique) iEEG channels with similar spectral signatures and 236 (99 unique) iEEG channels with dissimilar spectral signatures (all p < 0.05). Similar spectral signatures were distributed across medial frontal and parietal regions, as well as temporal and occipital regions (**Fig. 5A-B**).

**Figure 3.**
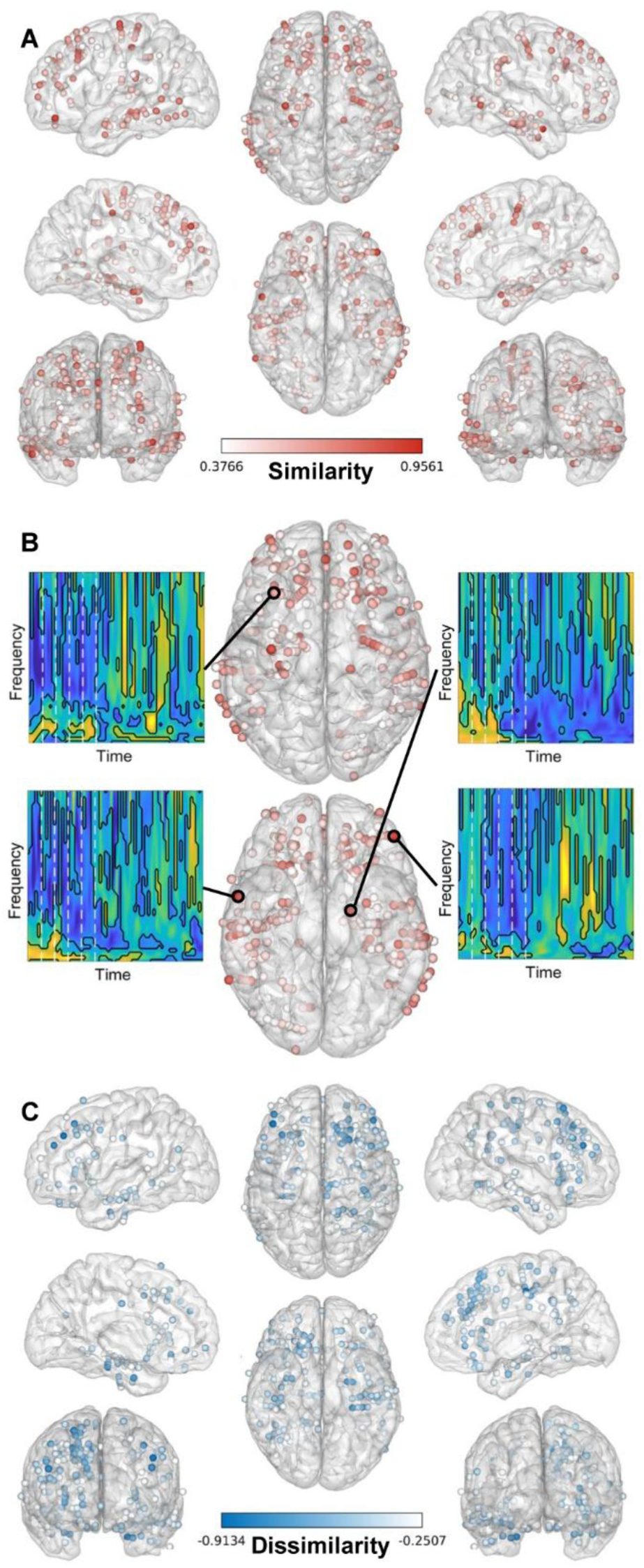
Intracranial EEG similarity and dissimilarity to scalp EEG FM ERSP effects. **A)** Spatial distribution of iEEG channels with spectral signatures significantly similar to the FM scalp ERSP effects in Fig. 2A (*z* > 1.96, *p* < 0.05; channels shown in red). Darker red indicates greater similarity. **B)** Time-frequency representations of t-statistics on power differences (test vs. sample) in four example channels selected by similarity index rank. The top left (rank 5/1,305) and bottom left (rank 13) channels are in expected medial and superior frontal regions. The top right (rank 4) and bottom right (rank 6) channels exhibit high similarity but are in inferior and lateral temporal regions. Black contours indicate significant clusters; electrode positions are shown on the brain. **C)** Spatial distribution of iEEG channels significantly dissimilar to the FM scalp EEG effect (*z* < −1.96, *p* < 0.05). Darker blue reflects greater dissimilarity.

**Figure 4.**
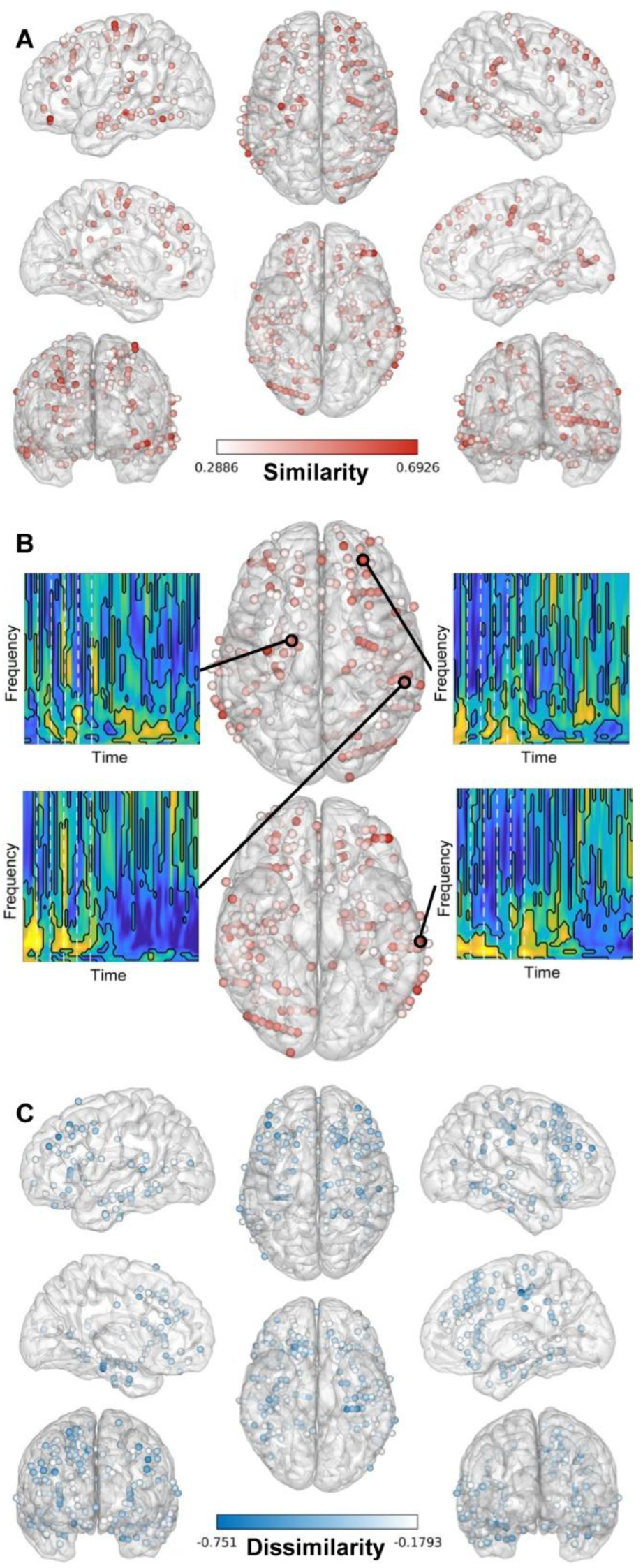
Intracranial EEG similarity and dissimilarity to scalp EEG bilateral posterior ERSP effects. **A)** Spatial distribution of iEEG channels with spectral signatures significantly similar to the bilateral posterior scalp ERSP effects in Fig. 2B (z > 1.96, p < 0.05; channels shown in red). Darker red reflects greater similarity. **B)** Time-frequency representations of t-statistics on power differences (test vs. sample) in four example channels selected by similarity index rank. The bottom left (rank 9/1,305) and bottom right (rank 2) channels are located in expected bilateral posterior regions. The top left (rank 3) and top right (rank 15) channels exhibit similarly high similarity but are situated in frontal and central areas. Black contours indicate significant clusters; electrode positions are shown on the brain. **C)** Spatial distribution of iEEG channels significantly dissimilar to the bilateral posterior scalp ERSP effects (z < −1.96, p < 0.05; channels shown in blue). Darker blue indicates greater dissimilarity.

**Figure 5.**
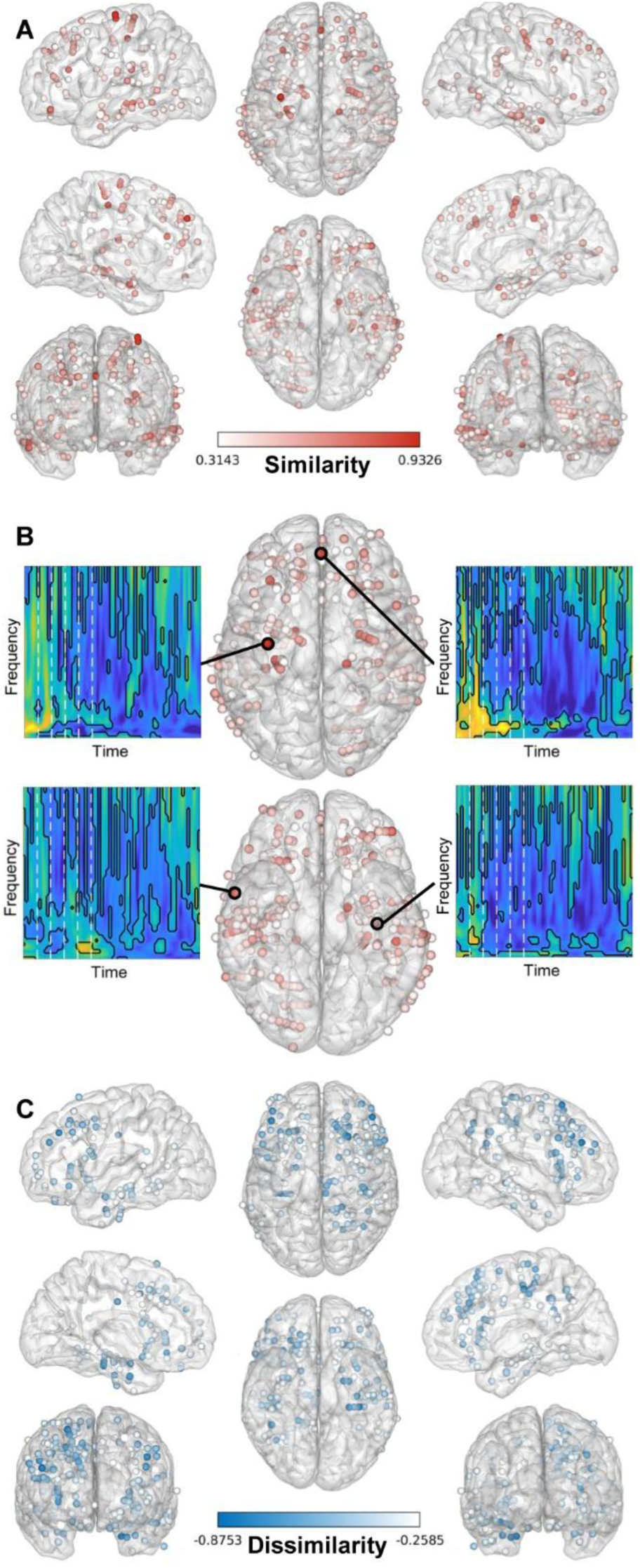
Intracranial EEG similarity and dissimilarity to scalp EEG central ERSP effects. **A)** Spatial distribution of iEEG channels with spectral signatures significantly similar to the central scalp ERSP effects in Fig. 2C (z > 1.96, p < 0.05; channels shown in red). Darker red indicates greater similarity. **B)** Time-frequency representations of t-statistics on power differences (test vs. sample) in four example channels selected by similarity index rank. The top left (rank 3/1,305) and top right (rank 7) channels are located in expected central regions. The bottom left (rank 4) and bottom right (rank 5) channels show similarly high similarity but are situated in inferior and lateral temporal areas. Black contours indicate significant clusters; electrode positions are shown on the brain. **C)** Spatial distribution of iEEG channels significantly dissimilar to the central scalp ERSP effects (z < −1.96, p < 0.05; channels shown in blue). Darker blue reflects greater dissimilarity.

Dissimilar spectral signatures were distributed across lateral PFC and lateral sensorimotor regions (**Fig. 5C**). Across all analyses, the same 177 channels (14%) were identified as similar, and the same 140 channels (11%) were identified as dissimilar.

These results demonstrate that distributed, overlapping iEEG spectral signatures of WM are similar to spatially non-overlapping scalp EEG signatures of WM. Results converge on expected effects based on scalp topographies, namely, likely medial frontal sources of FM and central scalp EEG effects, and likely inferior parietal, temporal, and occipital sources of bilateral posterior scalp EEG effects. However, they also reveal unexpected effects, namely, potential temporal and occipital sources of FM and central scalp EEG effects, and potential medial frontal sources of bilateral posterior scalp EEG effects. In all analyses, some similar and dissimilar iEEG spectral signatures were identified across neighboring channels within the same brain region, a common observation which may be attributed to the high spatial precision of iEEG (Mercier et al., 2022).

Lateral PFC regions were consistently identified as unlikely neuroanatomical sources of scalp EEG effects, consistent with the midline and posterior scalp topographies of the observed effects.

### Frontal midline theta hub signature of test sequence encoding

Having demonstrated ERSP signatures of WM distributed across the scalp topography linked to underlying iEEG in frontal, temporal, parietal, and occipital regions, we next investigated scalp EEG networks (i.e., top 35% PLV between each channel and all other channels (Johnson et al., 2022; Jones, Johnson, Tauxe, et al., 2020)). A significant cluster-corrected t-test revealed test > sample and sample > test hub effects that varied in time and frequency across the scalp topography (**Fig. 6A-C**). Sample > test effects were characterized by stimulus onset-locked phase-locking focused in the theta-alpha range (5-15 Hz) in central and bilateral posterior channels (p ≤ 0.048, Cohen’s d = 1.798). By contrast, test > sample effects were characterized by stimulus offset-locked phase-locking focused in the theta range (∼5 Hz) in FM channels (p = 0.036, Cohen’s d = 1.390), and in the theta-alpha range in central and bilateral posterior channels (p ≤ 0.030, Cohen’s d = 2.014). Additional sample > test effects were characterized by phase-locking in the theta-alpha range late during the delay in central channels (p ≤ 0.022, Cohen’s d = 0.812). These results demonstrate a FMT hub signature of test sequence encoding (see **Fig. 6A**), a bilateral posterior theta-alpha hub signature of temporally alternating sample and test sequence encoding (see **Fig. 6B**), and a central alpha hub signature of sample sequence maintenance (see **Fig. 6C**). A significant positive correlation between the FM hub signature of test sequence encoding and behavioral accuracy (ρ = 0.348, p = 0.041; **Fig. 6D**) revealed that individuals with superior performance had more densely connected FMT hubs during test sequence encoding.

**Figure 6.**
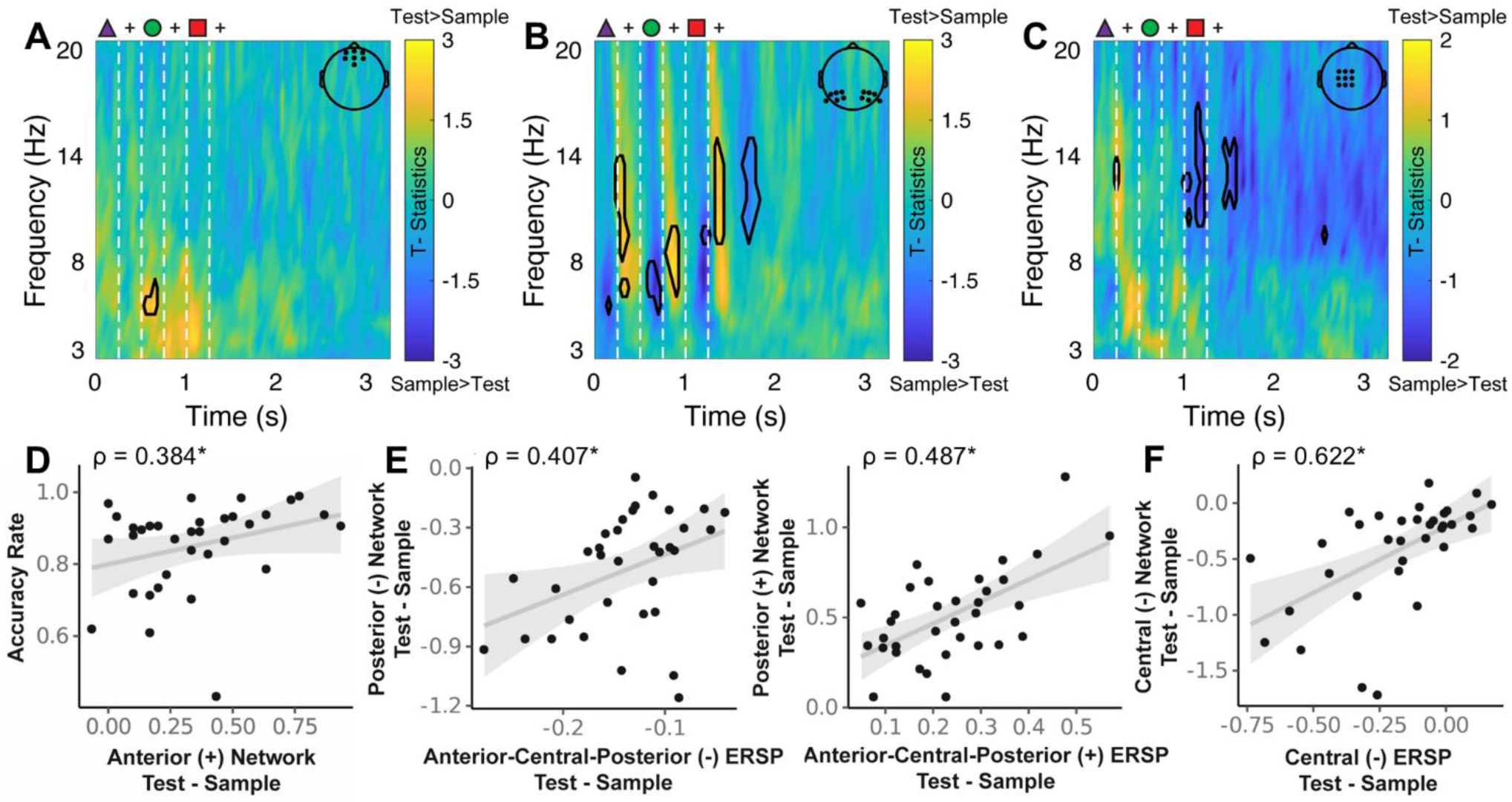
Task-related scalp EEG network hub effects. Time-frequency representations of t-statistics on network degree differences (test vs. sample) across FM, bilateral posterior, and central scalp EEG channels (insets). Time zero marks the onset of the first stimulus in each sequence. Geometric symbols indicate the three stimulus onsets (triangle, circle, square), and the plus sign denotes ISIs and delays. **A)** FM channels show increased theta phase-locking during test sequence encoding (yellow = test > sample; blue = sample > test). **B)** Bilateral posterior channels show increased theta-beta phase-locking during ISIs and reduced theta– beta phase-locking during stimulus presentation during test sequence encoding. **C)** Central channels show increased alpha phase-locking during test sequence encoding and sustained suppression during the delay. **D)** FM hub strength during test sequence encoding positively correlates with behavioral accuracy. Shading represents SEM. **E)** Bilateral posterior hub strength co-varies with ERSP effects across FM, bilateral posterior, and central regions: increased hub strength aligns with positive ERSP effects (test > sample), and decreased hub strength aligns with negative ERSP effects (sample > test). **F)** Central hub strength during the delay positively correlates with central ERSP effects reflecting sample maintenance.

Significant positive correlations between bilateral posterior hub and ERSP signatures (see **Fig. 2**) were observed for both the sample sequence (ρ = 0.407, p = 0.015) and test sequence encoding signatures (ρ = 0.487, p = 0.003; **Fig. 6E**), demonstrating that posterior hub connectivity and ERSPs co-occur during sample and test sequence encoding. A significant positive correlation between the central hub and ERSP signatures (see **Fig. 2C**) of sample sequence maintenance (ρ = 0.622, p < 0.001; **Fig. 6F**) revealed that central hub connectivity and ERSPs co-occur during sample sequence maintenance. No other correlations reached significance (p > 0.09). Taken together with ERSP similarity, results suggest that the bilateral posterior hub signature of temporally alternating sample and test sequence encoding (see **Fig. 6B**) reflects distributed inferior parietal, temporal, occipital, and medial frontal effects (see **Fig. 4A-B**). The central hub signature of sample sequence maintenance (see **Fig. 6C**) likely reflects distributed medial frontal and parietal, temporal, and occipital effects (see **Fig. 5A-B**).

### Frontoposterior theta synchrony signature of test sequence encoding

Having demonstrated FM and bilateral posterior ERSP and hub signatures of test sequence encoding, indicating broadly distributed effects as well as distinct rostral-superior frontal and inferior parietal sources, we investigated whether the hubs were in sync during test sequence encoding. Scalp EEG PLV data were averaged across FM-bilateral posterior channel pairs to create a single time-frequency representation of frontoposterior PLV. A significant cluster-corrected t-test revealed a test > sample effect characterized by stimulus offset-locked phase-locking in the theta-alpha range (2-12 Hz; p = 0.008, Cohen’s d = 1.025; **Fig. 7A**). A significant positive correlation between the PLV and ERSP signatures (see **Fig. 2**) of test sequence encoding (ρ = 0.387, p = 0.022; **Fig. 7B**) revealed that frontoposterior synchrony and ERSPs co-occur during test sequence encoding. No other correlations reached significance (p ≥ 0.13). Taken together with ERSP similarity, our results suggest that the frontoposterior PLV signature of test sequence encoding likely reflects medial frontal, temporal, and occipital effects, and synchrony between rostral-superior frontal and inferior parietal regions (see **Fig. 3-5**).

**Figure 7.**
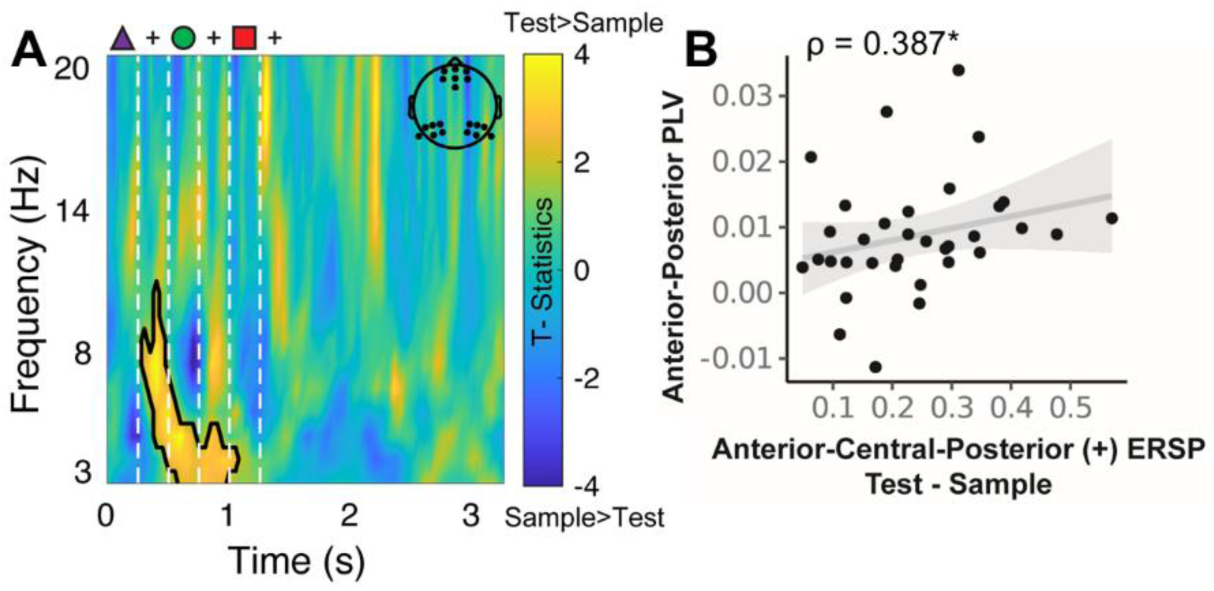
Task-related frontoposterior theta phase synchrony effects. Time-frequency representation of t-statistics on fronto-posterior PLV differences between test and sample sequences. Time zero marks the onset of the first stimulus in each sequence. White dashed lines indicate stimulus onsets and offsets. **A)** Frontoposterior channels exhibit increased theta phase synchrony during test sequence encoding (yellow = test > sample; blue = sample > test). Black contours outline significant clusters. **B)** Increased frontoposterior theta synchrony during test sequence encoding positively correlates with theta ERSP effects (test > sample), indicating that greater inter-regional coordination co-occurs with enhanced spectral activity. Shading represents SEM.

## Discussion

Our findings reveal how executive control in WM relies on localized spectral power changes and frequency-specific coordination across distributed brain networks. In our WM-DMS task, sequence comparison requires the engagement of executive processes to evaluate incoming stimuli against stored information and make goal-directed decisions. By contrasting test and sample sequences to isolate executive control processes, we observed increases in theta and alpha-beta power across FM and bilateral posterior regions during the test phase, accompanied by greater theta-alpha network hub strength and frontoposterior theta phase synchrony. In addition, individuals with higher WM accuracy exhibited denser connectivity among FMT hubs. Importantly, these dynamics were not confined to scalp-level patterns: using iEEG, we identified plausible anatomical generators of these signals, including medial frontal, parietal, temporal, and occipital regions. Together, these results suggest that executive control in WM is implemented through coordinated activity across local and large-scale systems, enabling the integration of stored and incoming information.

### Alternating theta and alpha-beta rhythms link regional activity to distributed networks

Our study demonstrates that FMT, posterior theta-beta, and central alpha-beta ERSPs were consistently associated with control processes during WM, revealing rapid alternations in their engagement and disengagement across task phases in scalp EEG. Specifically, FM channels exhibited increased test > sample theta power during ISIs, consistent with the recruitment of executive control during sequence evaluation (Cavanagh & Frank, 2014; Eisma et al., 2021; Itthipuripat et al., 2013). Bilateral posterior channels exhibited alternating theta-beta power increases during ISIs and decreases during stimulus presentation, supporting a dual role of posterior regions in maintaining internal representations during delays and modulating sensory input during comparison (Freedman & Ibos, 2018; Yang et al., 2017). These dynamics align with prior work suggesting that theta activity supports memory maintenance, while alpha and beta activity are involved in attentional gating and the suppression of irrelevant input (Fodor et al., 2020; Gutteling et al., 2022; Snyder & Foxe, 2010). Moreover, central channels showed similar test > sample theta power increases during ISIs but also displayed alpha-beta power reduction across the entire test sequence, consistent with prior work linking central beta reductions to sensory gating and memory content reactivation (Jensen & Mazaheri, 2010; Spitzer & Haegens, 2017). Together, these temporally specific and spatially distributed spectral effects point to a flexible coordination of sensory and executive systems during sequence comparison, enabling executive control and successful WM.

Network hubness, derived from PLV, exhibited similar task-phase dependent fluctuations as spectral power. Unlike spectral power, which reflects local signal amplitude, hub strength captures the extent to which a region is integrated with and functionally connected to other brain regions (Royer et al., 2022; Vecchio et al., 2017; Wijk et al., 2010). Bilateral posterior theta-alpha hubness increased during ISIs and decreased at stimulus onsets, mirroring power fluctuations across FM, posterior, and central channels. Moreover, theta PLV was significantly elevated between FM and bilateral posterior channels during test sequence encoding compared to sample sequence encoding. This result further supports theta as a key rhythm for inter-regional communication underlying executive control in WM, replicating prior findings emphasizing theta as a mechanism for large-scale cognitive coordination (Cohen & D’Esposito, 2016; Fiebelkorn & Kastner, 2019; Fries, 2015; Lisman & Jensen, 2013; Johnson et al., 2017, 2023; Cavanaugh & Frank, 2014; Parto Dezfouli et al., 2021).

Importantly, significant correlations between posterior hub strength and regional ERSP effects suggest that local theta-alpha activity supporting executive control is rooted in large-scale networks. A significant correlation was also found between theta PLV and theta-alpha ERSP effects, suggesting that long-range interactions work in tandem with local activity to support control in WM. These findings link regional spectral dynamics to broader network organization, bridging empirical EEG evidence with prior neural mass modeling work showing that changes in connectivity between neural populations can directly influence local power spectra (Zavaglia et al., 2008). Additionally, the FMT network hub was significantly associated with behavioral accuracy during WM, suggesting that FMT plays a key role in orchestrating cognitive control for successful task performance—a finding consistent with prior work linking FMT connectivity to executive function and behavioral outcomes (Fell & Axmacher, 2011; Cavanagh & Frank, 2014; Sauseng et al., 2005; Johnson et al., 2022; Gordon et al., 2018).

### Medial and posterior cortical origins of executive control signals

Given the observed links between regional and network-level spectral effects supporting control in WM, we asked where these effects originate. While scalp EEG offers excellent temporal resolution and consistent regional sampling across participants, spatial precision is compromised due to volume conduction. By contrast, iEEG provides both high temporal and spatial resolution, but its sparse and variable coverage across participants limits its use for group-level connectivity analyses (Parvizi & Kastner, 2018; Lagarde et al., 2022). To address both limitations, we examined scalp- and iEEG-derived ERSP effects as meaningful markers of spectral dynamics across modalities, with the goal of identifying plausible anatomical sources of scalp-derived executive control effects. Across all FM, central, and bilateral posterior scalp-defined ERSP effects, we found that similar iEEG spectral signatures were consistently localized to FM, parietal, temporal, and occipital regions, while dissimilar signatures tended to be concentrated in lateral PFC and sensorimotor areas.

Such widespread anatomical sources are not surprising given the observed ISI-related theta ERSP effects across FM, bilateral posterior, and central scalp channels. The prominence of iEEG effects in medial frontal regions is consistent with their well-established role in executive attention, goal maintenance, and cognitive control, particularly via FMT (Cavanagh & Frank, 2014; Ridderinkhof et al., 2004). This finding also aligns with prior scalp EEG studies identifying the anterior cingulate and pre-supplementary motor area, both core nodes in large-scale control networks, as dominant generators of FMT (Asada et al., 1999; Böttcher et al., 2025; Debener et al., 2005; Johnson et al., 2019; Schubert et al., 2025). Interestingly, similarity to scalp EEG effects was also observed in lateral and inferior temporal cortices, areas not typically implicated in control, but potentially engaged in contextual reactivation, semantic evaluation, or stimulus discrimination during sequence comparison (Axmacher et al., 2007, 2008; Herweg et al., 2020). These regions are obscured by the ears and not well captured in scalp EEG, highlighting the added value of intracranial data for mapping control-related processes beyond the canonical frontoparietal network.

In contrast, lateral PFC, often implicated in WM manipulation and top-down regulation (D’Esposito & Postle, 2015; Barbey et al., 2013; Johnson et al., 2017), tended to show dissimilarity in our scalp-iEEG comparisons. This pattern is consistent with prior iEEG work reporting that theta power in dorsolateral PFC decreases with increasing WM load and that such decreases predict superior behavioral performance (Brzezicka et al., 2019). While some studies have reported higher dorsolateral PFC theta during response inhibition (e.g., Khan et al., 2024), one specific form of executive control, the difference in task demands—with our sequence comparison emphasizing WM updating and selection more than response suppression—could partly account for the distinct theta patterns observed. Finally, we did not observe lateral frontal control signatures in the scalp EEG data, making our observation of dissimilarity in lateral PFC channels in the iEEG data unsurprising. Our FMT hub, linked to behavioral accuracy, may nonetheless be connected to nearby lateral PFC regions. Thus, the lateral PFC could be engaged in our task through low-frequency connectivity without prominent low-frequency activity, potentially accompanied by high-frequency activity, a dynamic not captured in our EEG analyses. Collectively, these findings point to dissociable medial and lateral PFC contributions to WM, with the lateral PFC’s role in executive function potentially involving neural mechanisms, such as high-frequency activity, that are more accessible through invasive approaches than scalp EEG.

### Limitations and future directions

Several limitations warrant consideration. First, while iEEG offers superior spatial resolution, its coverage is constrained by clinical needs for seizure localization and varies across patients, restricting consistent sampling across individuals (Parvizi & Kastner, 2018). While some iEEG studies have applied graph-theoretical approaches to study connectivity directly, these are typically constrained to limited subnetworks with sufficient coverage (Lagarde et al., 2022). Due to heterogeneous electrode placement and sparse spatial sampling within individuals, we focused on regional ERSP analysis rather than network metrics such as PLV or graph topology. This choice limits the conclusions that can be drawn about dynamic connectivity in iEEG, though our ERSP-based similarity approach offers a reasonable proxy given the significant correlation between the control-related ERSP and PLV effects. Second, our task was designed to isolate sequence comparison under controlled conditions, limiting generalizability to more naturalistic or complex WM demands. Third, given the observational nature of EEG and iEEG, our study cannot establish causal relationships between neural dynamics and behavior.

Despite these limitations, our approach offers a key methodological strength. Scalp EEG source localization faces inherent limitations, as the inverse problem does not yield a unique solution without additional assumptions or constraints (Grech et al., 2008; Michel & Brunet, 2019; Lai et al., 2018). Our strategy of linking scalp EEG-derived ERSP effects to iEEG-derived ERSP effects provides a novel solution that is increasingly recognized as robust (Seeber et al., 2019; Subramanian et al., 2025). By leveraging iEEG to constrain and contextualize scalp EEG findings, we offer a cross-modal framework that advances source interpretation, with a proof of concept in the study of cognitive control. Future research may improve upon this approach by incorporating simultaneous scalp and iEEG recordings to directly map surface signals to their intracranial sources within individuals and analyze connectivity through coordinated multi-center datasets that afford participant selection based on spatial coverage of multiple target regions. Future studies may also evaluate the generalizability of our results to more naturalistic tasks, clinical populations with executive dysfunction, and experimental paradigms using brain stimulation to test causal mechanisms (Cohen, 2017).

High-density electrode arrays, improved source modeling techniques, and unified behavioral designs will be essential for resolving how local and distributed oscillatory dynamics jointly support executive control in the human brain.

## Acknowledgements

We thank Vasanth Kommu for assistance with scalp EEG data collection, and Drs. Jack Lin, Peter Weber, Kenneth Laxer, and Edward Chang for assistance with intracranial EEG data collection. This research was supported in part through the computational resources and staff contributions provided for the Quest high-performance computing facility at Northwestern University, which is jointly supported by the Office of the Provost, the Office for Research, and Northwestern University Information Technology. The study resulting in this publication was assisted by a grant from the Undergraduate Research Grant Program, which is administered by Northwestern University’s Office of Undergraduate Research. However, the conclusions, opinions, and other statements in this publication are the author’s and not necessarily those of the sponsoring institution.

